# The *Cryptococcus neoformans* Titan cell is an inducible and regulated morphotype underlying pathogenesis

**DOI:** 10.1101/190587

**Authors:** Ivy M. Dambuza, Thomas Drake, Ambre Chapuis, Leanne Taylor-Smith, Nathalie LeGrave, Tim Rasmussen, Matthew C. Fisher, Tihana Bicanic, Thomas S. Harrison, Marcel Jaspars, Robin C. May, Gordon D. Brown, Raif Yuecel, Donna M. MacCallum, Elizabeth R. Ballou

**Affiliations:** Institute of Medical Science, University of Aberdeen, Aberdeen, UK; Institute of Microbiology and Infection, School of Biosciences, University of Birmingham, Birmingham, UK; Marine Biodiscovery Centre, Department of Chemistry, University of Aberdeen, Aberdeen, UK; Francis Crick Institute, London, UK; Institut für Biochemie, Universität Würzburg, Wurzburg, Germany; Department of Infectious Disease Epidemiology, Imperial College London. London, UK; Institute of Infection and Immunity, St George’s University of London, London, UK

## Abstract

Fungi undergo changes in cell shape in response to environmental stimuli that drive pathogenesis and niche adaptation, such as the yeast-to-hyphal transition of dimorphic fungi in response to changing temperature. The basidiomycete *Cryptococcus neoformans* undergoes an unusual morphogenetic transition in the host lung from haploid yeast to large, highly polyploid cells termed Titan cells. Titan cells influence fungal interaction with host cells, including through increased drug resistance, altered cell size, and altered Pathogen Associated Molecular Pattern exposure. Despite the important role these cells play in pathogenesis, understanding the environmental stimuli that drive the morphological transition, and the molecular mechanisms underlying their unique biology, has been hampered by the lack of a reproducible *in vitro* induction system. Here we demonstrate reproducible *in vitro* Titan cell induction in response to environmental stimuli consistent with the host lung. *In vitro* Titan cells exhibit all the properties of *in vivo* generated Titan cells, the current gold standard, including altered capsule, cell wall, size, high mother cell ploidy, and aneuploid progeny. We identify bacterial peptidoglycan as a serum compound associated with shift in cell size and ploidy, and demonstrate the capacity of bronchial lavage fluid and *E. coli* co-culture to induce Titanisation. Additionally, we demonstrate the capacity of our assay to identify established and previously undescribed regulators of Titanisation *in vitro* and investigate the Titanisation capacity of clinical isolates and their impact on disease outcome. Together, these findings provide new insight into the environmental stimuli and molecular mechanisms underlying the yeast-to-titan transition and establish an essential *in vitro* model for the future characterization of this important morphotype.

**Author Summary:** Changes in cell shape underlie fungal pathogenesis by allowing immune evasion and dissemination. *Aspergillus* and *Candida albicans* hyphae drive tissue penetration. *Histoplasma capsulatum* and *C. albicans* yeast growth allows evasion and dissemination. As major virulence determinates, morphogenic transitions are extensively studied in animal models and *in vitro*. The pathogenic fungus *Cryptococcus neoformans* is a budding yeast that, in the host lung, switches to an unusual morphotype termed the Titan cell. Titans are large, polyploid, have altered cell wall and capsule, and produce haploid daughters. Their size prevents engulfment by phagocytes, yet they are linked to dissemination and altered immune response. Despite their important influence on disease, replicating the yeast-to-Titan switch *in vitro* has proved challenging. Here we show that Titans are induced by host-relevant stimuli, including serum and bronchio-alveolar lavage fluid. We identify bacterial peptidoglycan as a relevant inducing compound and predict an *in vivo* Titan defect for a clinical isolate. Genes regulating *in vivo* Titanisation also influence *in vitro* formation. Titanisation is a conserved morphogenic switch across the *C. neoformans* species complex. Together, we show that Titan cells are a regulated morphotype analogous to the yeast-to-hyphal transition and establish new ways to study Titans outside the host lung.

## Introduction

Fungi change shape in response to environmental stimuli. These morphogenic transitions drive pathogenesis and allow fungi to occupy different environmental niches. Dimorphic fungi undergo a yeast-to-hyphal transition in response to changing temperature, and the pleomorphic gut resident fungus *Candida albicans* integrates diverse signals depending on its local environment [1, 2]. The basidiomycete *Cryptococcus neoformans* undergoes an unusual transition in the host lung from haploid yeast-phase growth to apolar expansion and endo-reduplication, producing large, highly polyploid cells termed Titan cells [3, 4]. While there is growing evidence of the important role Titan cells play in disease [5-8], understanding the mechanisms underlying the yeast-to-Titan transition remains challenging due to the lack of an *in vitro* model.

C. *neoformans* is an environmental human pathogen that causes cryptococcal meningitis when inhaled yeast and spores disseminate to the central nervous system and brain. The fungus infects an estimated 1 million people worldwide each year and is responsible for between 140,000 and 600,000 deaths, primarily in sub-Saharan Africa [9-11]. Although the majority of patients are immunocompromised, a growing number of infections are seen in immuno-competent individuals [12-14]. Long term azole therapy is associated with relapse due to drug resistance and the emergence of heteroresistance [14-16]. *C. neoformans* grows preferentially as an encapsulated budding yeast under physiologically relevant conditions and during culture in standard microbial media, and the vast majority of research has focused on the yeast form. However, there are early clinical reports of Titan cells, in which large encapsulated yeast were isolated from the lung and brain of infected patients [17, 18]. In both cases, cell size was dependent on growth condition, shifting from >40 μm in patient samples to <20 μm during in vitro culture and back to >40 μm in murine infection. Cruickshank et al. also report distinct capsule and cell wall structure of enlarged cells[18]. Despite this clear morphological transition, both early reports concluded that the patient samples represented atypical isolates.

However, far from being unusual outliers, it is now clear that Titan cells represent a unique aspect of cryptococcal biology. Recent work in mouse models of infection have demonstrated that Titan cells comprise 20% of fungal cells in the lung and are associated with dissemination to the brain and a non-protective immune response [5, 7, 8, 19]. Titanisation requires the activity of the Gα protein Gpa1 and the G protein coupled receptor Gpr5, as well as the mating pheromone receptor Ste3a, likely targeting the cAMP/PKA pathway [20]. Transcription factors that influence cAMP-regulated capsule and melanin also influence Titanisation [20-22]. However, the environmental triggers of Titanisation remain unknown, and reports of *in vitro* Titanisation have not led to a robust *in vitro* protocol for their generation [20, 23-25].

The analysis of Titan cells recovered from infected mice has led to the identification of four defining features: Titans are larger than 10 μm, are polyploid (typically 4-8C, although higher ploidies have been reported), have a tightly compacted capsule, and have a dramatically thicker cell wall [6, 8, 18]. Here we report a simple and robust protocol for the *in vitro* generation of cells matching this definition. We validate the capacity of our protocol to identify genes required for Titanisation, and predict the capacity of clinical isolates to form Titans. Finally, we identify environmentally relevant ligands that trigger the yeast-to-Titan transition and begin to dissect the underlying molecular mechanisms that drive this novel virulence mechanism.

## Results

### Large polyploid cells can be induced *in vitro*

While investigating the impact of nutrient starvation on virulence factor production, we observed that when *C. neoformans* cells grown in Yeast Nitrogen Base (YNB) with 2% glucose were transferred to 10% HI-FCS at 5% CO_2_, 37°C, large cells (up to 50 μm) formed after several days (Fig. 1A, 3 days; Fig. S1A, 7 days). These cells expressed a compact capsule that was readily distinguished from the more typical yeast capsule by India ink staining (Fig. 1A). Similar effects were observed for cells grown in 10% native FCS or heat inactivated (HI)-FCS or but not for culture-matched cells transferred to 1xPBS. Given the reported capacity of *C. neoformans* to form polyploid Titan cells, these large cells were examined for DNA content. Induced cells were passed through an 11 μm filter to enrich for large cells. Both fractions were collected, fixed and stained for DNA content (Fig. 1B,C, S1B gating strategy). While un-induced cells showed distinct 1C and 2C peaks representing progression of haploid yeast through the cell cycle, induced cells additionally showed discrete peaks consistent with populations of higher ploidy cells. The filtered population was enriched for large cells with increased DNA content (Fig 1C), consistent with these large cells being polyploid Titan cells.

**Figure 1:**
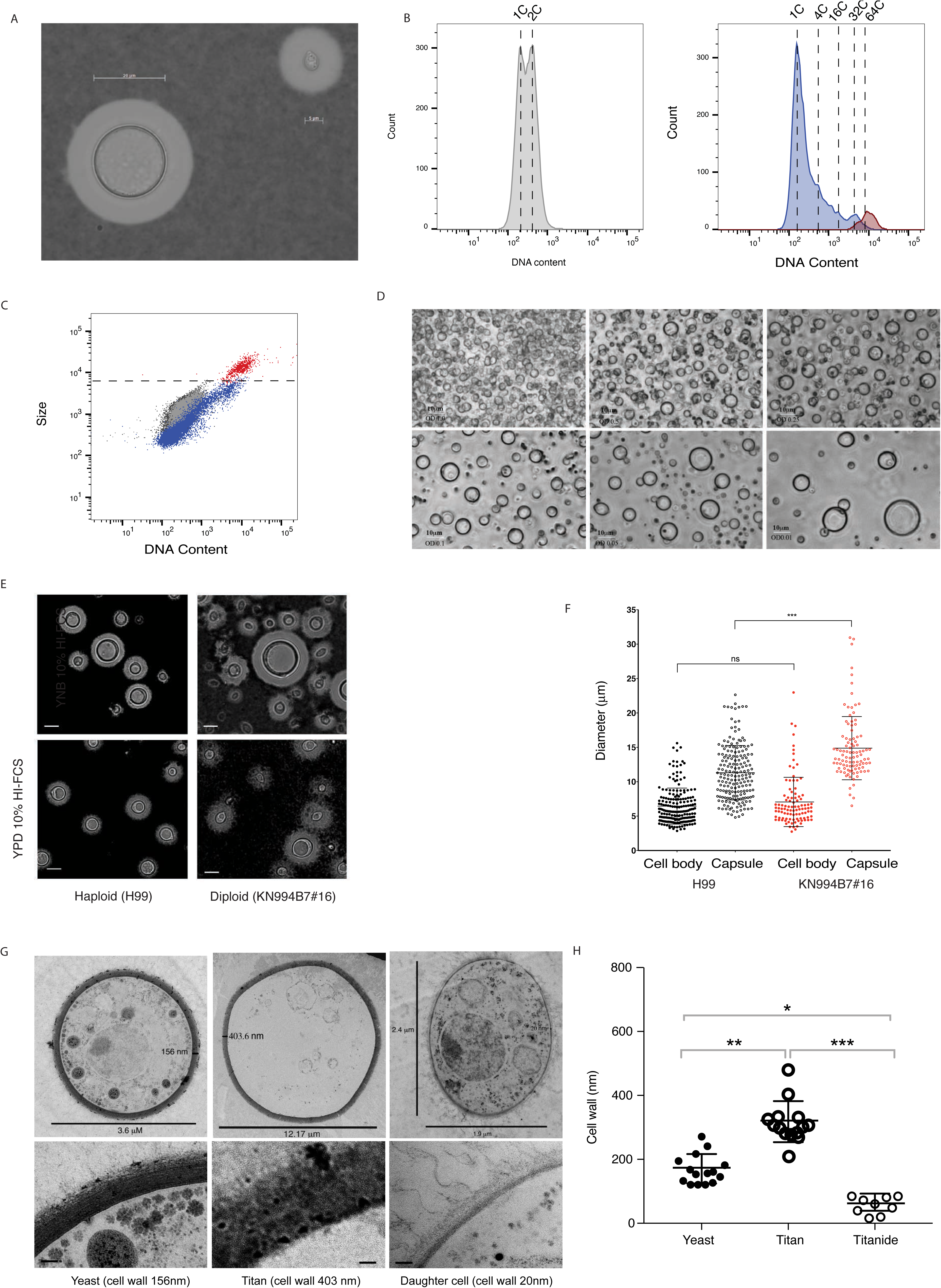
Titan cells can be induced *in vitro* and have all the properties of *in vivo* induced cells. H99 haploid cells pre-grown in YNB and incubated in 10%HI-FCS at 37°C, 5%CO_2_ for 3 days. Resulting colonies were A) counterstained with India ink to reveal capsule and B) fixed and stained for DNA content relative to haploid and diploid controls. Induced cells were passaged through an 11 μm filter to enrich for large cells. C) H99 haploid cells were pre-grown in YNB and incubated in 10%HI-FCS at the indicated OD_600_ at 37°C, 5%CO_2_ for 3 days. Scale bar = 10 μm. D,E) H99 haploid and KN994B7#16 diploid cells pre-grown in either YNB or YPD were inoculated into 10%HI-FCS at OD_600_=0.001, (37°C, 5%CO_2_, 3 days) and then counterstained with India ink and analysed. Representative micrographs (D) and diameters for >200 cells excluding or including capsule (E) are shown. Significance was assessed using Mann Whitney U (***p<0.0001). F) *In vitro* induced cells were analysed by TEM. Representative micrographs (F) and cell wall thickness for Yeast (n=15), Titan (n=14), and Titanide (n=9) cells (G). Significance was assessed using Kruskal-Wallis and Dunn's multiple comparisons tests (* p=0.027; ** p=0.002; ***p<0.0001).

Titan cells observed *in vivo* typically comprise up to 20% of the total cell population, and lower inocula are associated with an increase in the proportion of Titan cells [20]. We likewise observed that a minority of cells fell into the >10 μm category and that the percentage and overall cell size increased at decreasing inoculum concentration (Fig. 1D). While large cells were readily observed at OD_600_=0.25, there was an increase in the frequency and size of larger cells at lower optical densities (OD_600_ 0.05, 0.01). Optimization revealed that larger than average cells (10-12 μm) could be observed after 24 hr, but that cells approaching 15 μm were readily observed after 72 hr. Therefore, all subsequent characterizations were performed using an overnight culture of YNB+Glucose grown cells inoculated into 1xPBS + 10% HI-FCS at OD_600_ = 0.001 and incubated for 72 hr at 37°C, 5% CO_2_. The induced phenotype was reproducible across labs and users (University of Aberdeen (TD, ERB), University of Birmingham (ERB, LTS)).

### *In vitro* generated large cells are Titan cells

Where yeast cells typically range in size from 5-7 μm, Titan cells are defined as being >10 μm (cell body) or >15 μm (including capsule) [24]. When we measured the cell body diameter of induced cells, we observed a size range spanning 3-15 μm (Figure 1E,F). Cells >10 μm represented 15.72 ± 4.46 % of the population. When capsule was taken into account, the proportion of cells >15 μm increased to 30% (Fig. 1F). Cell size and ploidy are proportional, and we tested the impact of base ploidy on induced cell size. When cell body alone was considered, there was a shift in the size and frequency of cells >10 μm between haploid (H99) and diploid (KN994B7#16) base ploidy (Fig. 1F p=0.0785). When capsule was taken into account, the difference in size became highly significant (Fig. 1F; p<0.0001), suggesting that capsule size increases with ploidy. However, within the induced H99 population, cell:capsule ratios were not uniform across the entire population (Figure S1B). Cell body to capsule ratio was significantly smaller for cells >10μm (1.425 vs. 1.696; p<0.0001). This is consistent with reports that the capsule of Titan cells is more densely crosslinked than that of non-Titan yeast[24].

Titan cells have thicker cell walls than yeast cells and contain a single large vacuole of unknown function. TEM analysis (Fig. 1 G,H) revealed that cells >10 μm had significantly thicker cell walls (314.7 ± 64.0 nm) than those 10< μm (167.3 ± 46.2 nm; p = 0.002). Large cells were mostly devoid of organelles. In rare instances, some cytoplasmic material could be observed along the cell cortex, consistent with the presence of a large vacuole (Fig. S1D). A third population of small (2-4 μm) cells was also observed (Fig. 1 G). These small, encapsulated cells resembled yeast in that they appeared metabolically active, with ribosomes, mitochondria, and nucleus readily visualized, and capsule observed extending from the cell wall. However, where yeast and yeast daughters are round, these tended to be oval and had significantly thinner cell walls (56.14 ± 26.8 nm; p=0.026) (Fig 1H). These cells appear to be distinct from the previously reported microcells, defined as <1 μm [26], and we are not able to find reports of similar small ovoid cells in other TEM analyses of *C. neoformans* nor have we observed these cells in other TEM analyses we ourselves have conducted under non-Titan inducing conditions. We speculate that these cells are newly generated Titan daughter cells, which we here term Titanides, in order to distinguish them from yeast daughter cells.

Titan cells are uninucleate, polyploid, and produce haploid or aneuploid daughters [6, 24]. The large induced cells likewise contained a single nucleus, and were capable of passing DNA to daughter cells (Fig. 2A). To study these daughters, individual large cells were isolated by microdissection and allowed to proliferate on YPD agar at 30°C for 16 hr. Individual daughters were dissected and further incubated on YPD agar at 30°C for 24 hr. Figure 2B shows representative flow cytometry data measuring DNA content for fourteen daughter cells arising from a single Titan mother. These daughters were diploid or aneuploid relative to the H99 haploid parent, consistent with the environmentally-induced change in cell ploidy characteristic of Titan cells. While some daughters resolved back to haploidy within 1 month (25°C, YPD agar), others were stable within this time scale. Based on these data, including size, capsule, cell wall, and ploidy, we suggest that YNB-serum induced large cells are in fact *bona fide* Titan cells.

**Figure 2:**
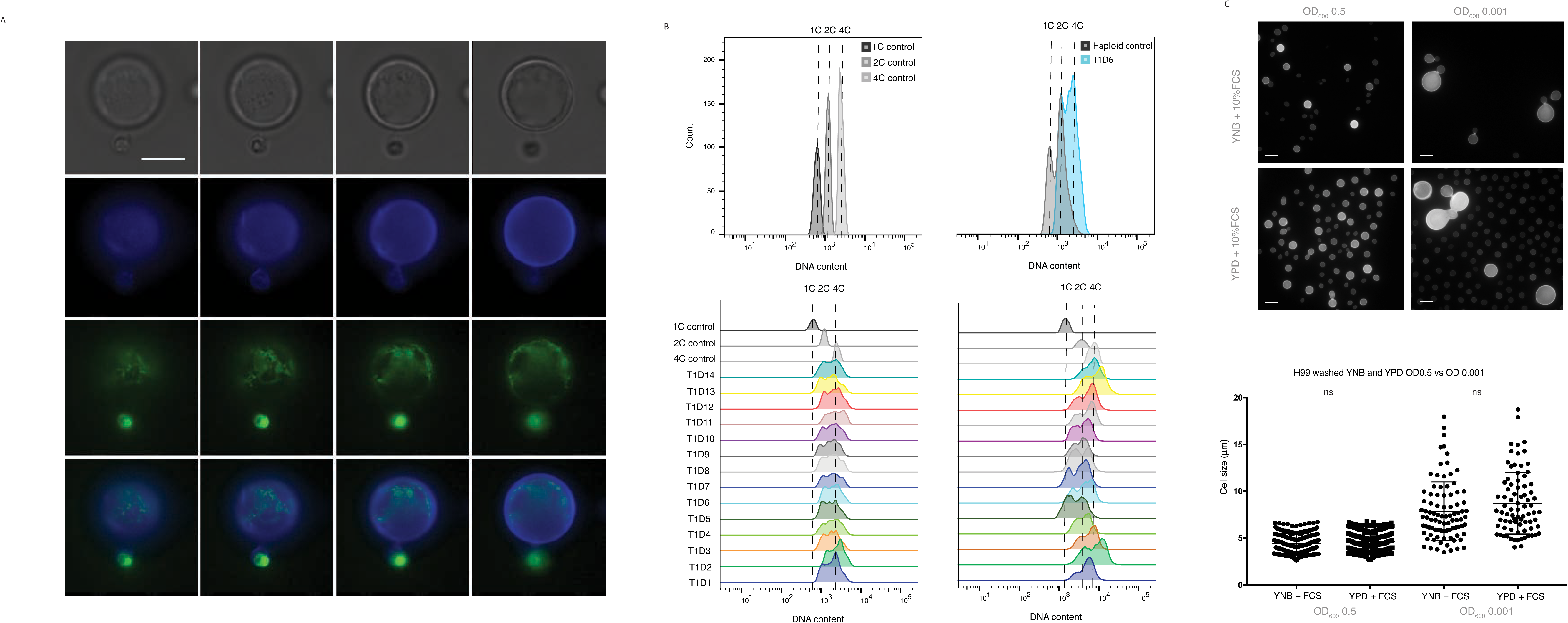
Daughter cells from *in vitro* Titans have altered ploidy relative to the haploid parent. A) Representative budding Titan cell. H99 haploid cells were induced and stained for chitin (CFW, 10 μg/ml) and DNA (SytoxGreen, 50 μm/ml). Scale bar = 5 μm, Z=2 μm sections. B) Daughters arising from a single Titan mother were isolated and allowed to proliferate for 72 hr at 30°C on YPD agar prior to fixation and staining for DNA content (DAPI). Populations were analysed by flow cytometry. (top) Control gates (1C, 2C, 4C) are shown aligned to representative overlays of ungated haploid control and Titan Daughter #5 (T1D5) populations. (bottom) DNA content of 14 independent daughter colonies 48 hr after isolation (right) and again after 1 month incubation at RT on YPD agar. C) H99 cells pre-cultured in either YPD or YNB were washed 6 times in 1xPBS and then inoculated at the indicated OD in 10% FCS (3 days, 37°C, 5%CO_2_). Representative micrographs and cell size quantification are shown.

### Titan cells form in response to sequential signals

Our *in vitro* induction protocol is a two-step process: cells are first incubated under minimal media conditions, and then induced to undergo the yeast-to-Titan switch via exposure to FCS. Given the observed impact of inoculation density on Titanisation (Fig. 1D), we tested whether changes in secreted factors dependent on cell density might repress Titanisation by cells grown in rich media. YPD-grown cells were washed 6 times in PBS to remove residual exogenous compounds and incubated in 10% HI-FCS at OD_600_=0.5 or 0.001 (Fig 2C). Titan cells were not observed in either YPD or YNB-pre-grown cultures at OD_600_=0.5. At OD_600_=0.001, washed YPD-grown cells produced large cells at rates similar to YNB-grown cells (p>0.99). However where YNB-grown Titan cells produce disproportionately small daughters, similar in size to yeast daughters, YPD-grown large cells frequently produced large buds, proportional to the large mother cell. YPD-grown mother-daughter pairs also tended to be dysmorphic, with defects in cytokinesis (Fig. 2C). These data suggest that cell density influences the generation of large cells, but density-dependent secreted factors alone are not sufficient to explain the yeast-to-Titan transition.

### Titanisation can be induced by bacterial cell wall components

Titan cells have been identified in the host lung and brain, but have not been observed circulating in the blood or CNS. To test the impact of host-relevant inducing compounds, we asked whether murine Bronchial Alveolar Lavage (BAL) extract could induce Titan cells. When 10% BAL was used in place of FCS, we observed large polyploid cells similar to FCS-induced Titans (Fig. 3A). Daughter cells arising from BAL-induced Titans were microdissected and cultured as described above for FCS-induced daughters. BAL-induced Titan daughters also exhibited a shift in base ploidy to 2C and 4C, with daughters arising from the same mother showing a range in base ploidy (Fig. 3B). Therefore, BAL fluid and FCS share the same capacity to induce the yeast-to-Titan transition.

**Figure 3:**
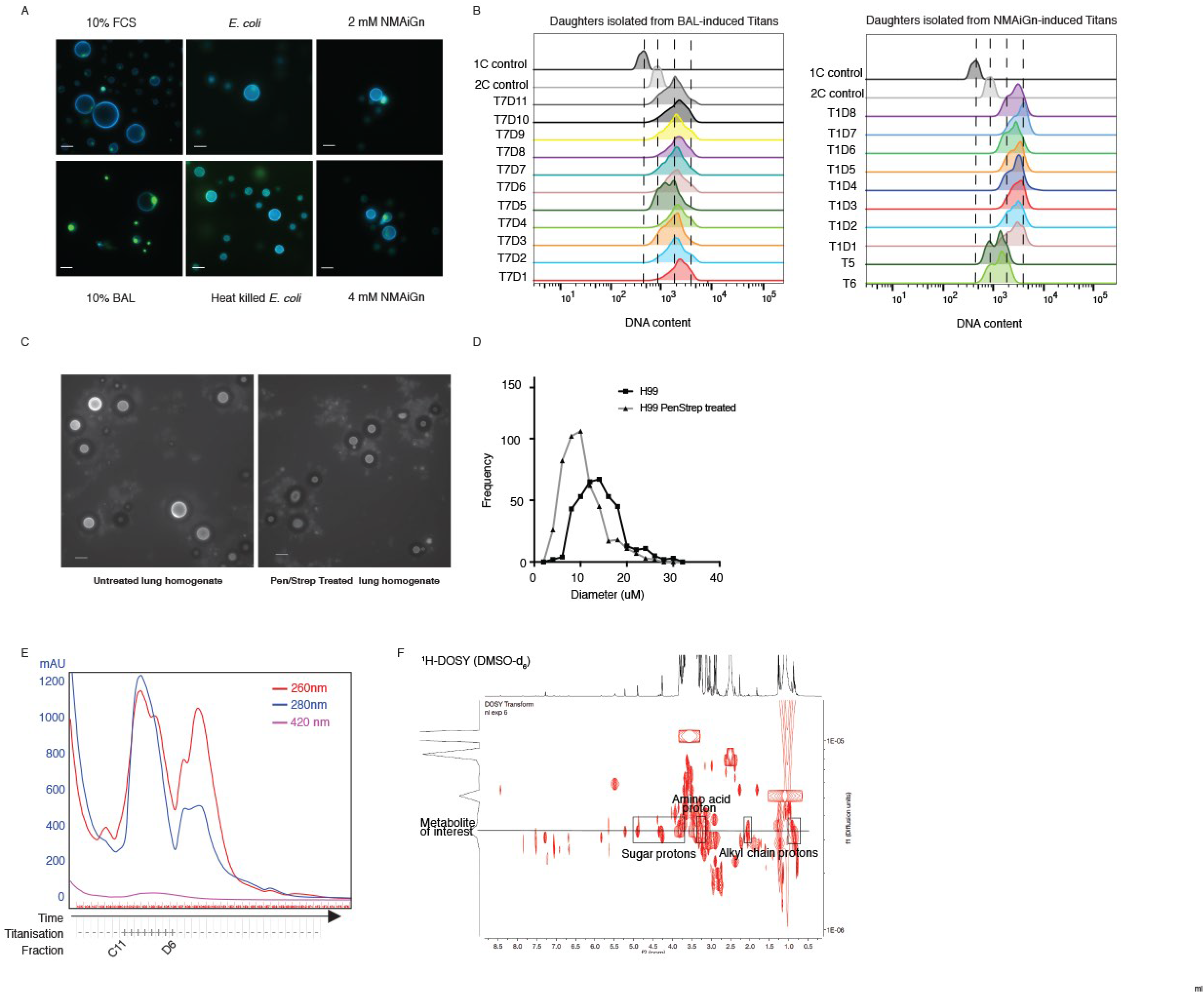
Titan cell induction is mediated by host relevant ligands and pre-culture condition. A) H99 cells pre-cultured in YNB were induced for Titans following 3 days growth in 10% FCS, 15% BAL, or 10 μM or 20 μM N-Acetylmuramyl-L-alanyl-D-isoglutamine (NMAiGn) as indicated. Representative micrographs are shown for live cells (CFW, SytoxGreen). Scale bar = 10 μm. B) Daughters arising from Titan mothers induced using BAL (left, T7) or NMAiGn (right, T1, T5, T6) were isolated and allowed to proliferate for 72 hr at 30°C on YPD agar prior to fixation and staining for DNA content (DAPI). Populations were analysed by flow cytometry. C,D) Lung homogenates from mice unexposed or exposed to Pen/Strep for 7 days prior to *C. neoformans* inhalation infection. Representative micrographs (CFW, scale bar = 20 μm) (C) and quantification (D) are shown. n>300 for each condition. E,F) FCS was fractionated by size exclusion chromatography (E) and the resulting fractions were tested for capacity to induce Titan cells. F) Analysis by ^1^H NMR and ^1^H-^13^C HSQC revealed peaks consistent with sugar and amino acid structures.

BAL extract is predicted to include lung-resident bacteria, a normal component of the host microbiome. To model the role of the host microbiome on Titanisation, we tested *C. neoformans* and *Escherichia coli* co-cultures for Titan induction. Co-culture of YNB-grown *C. neoformans* and either live or heat-killed *E. coli* was sufficient to induce Titan cells after 48 hr (Fig 3A). We tested the *in vivo* relevance of the host microbiome on Titan cell induction by comparing fungal cell size in the lungs of infected mice to fungal cell size in the lungs of mice pre-treated with antibiotic water for 7 days. There was no difference in fungal CFUs between treated and untreated mice, whereas bacteria could be cultured from the lungs of untreated mice, but not treated mice. We examined lung homogenates (Fig 3C) and histology (Fig. S2A) for evidence of Titanisation. Although large cells were observed in homogenates from both treated and untreated mice, there was a significant reduction in maximum cell size for treated mice (Fig 3C, (p<0.0001)), suggesting that clearance of lung bacteria reduced the degree of Titanisation in the lungs. Exposure of *C. neoformans* to antibiotic had no impact on Titanisation *in vitro* (data not shown).

Together, these data suggest that host-relevant ligands in both FCS and BAL modulate *C. neoformans* Titanisation and demonstrate that bacterial factors influence *C. neoformans* morphogenesis. We therefore aimed to determine the minimum components of FCS necessary to trigger large cells. HI-FCS was fractionated by size exclusion chromatography, and YNB-grown H99 cells were incubated in 10% compositions of each fraction in 1xPBS. Large cells (>10 μm) were observed in cultures incubated with fractions from wells C11-D11, matching a large peak that eluted after 11 min (Fig. 3E, S2B). Comparison to size standards suggested that compounds in this peak are in the range of 500 Daltons. We further fractionated the pooled sample by HPLC and tested the fractions for inducing activity. Analysis by ^1^H-NMR and DOSY suggested a complex mixture of at least 9 different compounds (Fig. 3F). NMR data suggested the presence of a metabolite with a sugar component (δ_H_ 4.89, 4.32/4.30, 4.27, 3.79 and 3.70 ppm) and an alkyl chain (δ_H_ 2.08, 2.03, 0.87ppm). Additionally, ^1^H NMR and ^1^H-^13^C HSQC experiments exhibited a methylene (δ_H_ 3.12 ppm, 52.4 ppm) likely to be located in alpha conformation to a carbonyl and an amino group, which suggested the presence of an amino acid substructure in this metabolite. These features are consistent with peptidoglycan structures. Coupled with our observation that bacterial cell wall is capable of triggering Titanisation, we hypothesized that this metabolite might represent a bacterial peptidoglycan.

Muramyl tetrapeptides (MTP) are peptidoglycan subunits common to the cell walls of Gram negative, Gram positive, and myco-bacteria. Muramyl tetrapeptides consist of an ether of N-acetylglucosamine (GlcNAc) and lactic acid (MurNAc), plus a species-specific tetrapeptide. MTPs have been shown to act as signaling molecules in both mammalian and fungal systems through interaction with Leucine Rich Repeat (LRR) domains on target proteins, including mammalian Nod receptors and *C. albicans* adenylyl cyclase [27]. MTP and its derivatives were identified as potent inducers of the yeast-to-hyphal transition in *C. albicans* following spectroscopic analysis of serum, which was shown to contain low levels of bacterial cell wall component [27]. Synthetic Muramyl Dipeptide, *N*-Acetylmuramyl-L-alanyl-D-isoglutamine (NMAiGn), is structurally similar but not identical to MTP. ^1^H NMR analysis of NMAiGn was consistent with the peptidoglycan peaks identified in the FCS fractions. To confirm the role of MTP derivatives in *C. neoformans* morphogenesis, Titan cells were induced using 2 mM or 4 mM NMAiGn (the concentration sufficient to trigger the yeast-to-hyphal switch in *C. albicans*). Cells incubated with NMAiGn exhibited limited proliferation; however, cells >10 μm were present at both concentrations, consistent with a yeast-to-Titan switch (Fig. 3A). Individual large cells were isolated by microdissection and allowed to proliferate for 17 hr at 30°C on YPD agar. Of 6 large cells isolated, all 6 proliferated to form colonies. For four of these colonies, individual daughters, distinguishable through their reduced size relative to the mother, were again isolated and allowed to proliferate for a further 72 hrs. The remaining 2 colonies (T5, T6) from the original large cells were analysed in aggregate. Each of the resulting lineages was analysed by flow cytometry for cell size and ploidy. In NMAiGn-induced daughter cells, we observed an overall increase in ploidy, with the majority of colonies arising from individual daughters having a 4C base ploidy. A representative lineage is shown in Figure 3B. Aggregate samples were more heterogeneous and included 2C and 4C cells consistent with diploid daughter lineages. Daughter cells arising from different mothers were more similar to their siblings, consistent with the shift in ploidy arising as a consequence of events during induction, rather than following isolation of the mother cell onto YPD agar. Together, these data demonstrate a role for peptidoglycan such as MTP during *in vitro* Titanisation and suggest that bacterial components influence Titanisation *in vivo*.

### The cAMP pathway positively regulates *in vitro* Titanisation

Muramyl dipeptide is thought to interact directly with the LLR domain of adenylyl cyclase, and the cAMP signal transduction cascade is believed to regulate Titanisation *in vivo* [20, 22, 27]. However, addition of exogenous cAMP at levels sufficient to induce capsule failed to induce Titan cells in either YPD or YNB-grown cultures (data not shown). The avirulence of mutants deficient in cAMP signal transduction has precluded direct testing of this model [20, 28, 29]. We therefore examined the influence of *GPA1, CAC1*, and *PKA1*, as well as *RIC8*, a Gα Guanine Nucleotide Exchange Factor (GEF) for Gpa1, on *in vitro* Titanisation [30]. Strains deficient in each of these genes failed to generate large cells in our assay (Fig. 4A,B). The G-protein coupled receptor Gpr5 is required for Titanisation, and the *gpr4Δgpr5Δ* strain exhibits a significant reduction in Titan cell production *in vivo* [7, 20]. We likewise observed a decrease in the frequency of Titan cells *in vitro* in the *gpr4Δgpr5Δ* strain (Fig. 4A,B). Consistent with the incomplete defect observed *in vivo[20]*, rare Titan cells could be observed *in vitro* for this strain (Fig. 4B). Together, these data demonstrate that *in vitro*-induced Titan cells are regulated via a similar pathway to *in vivo* Titan cells.

**Figure 4:**
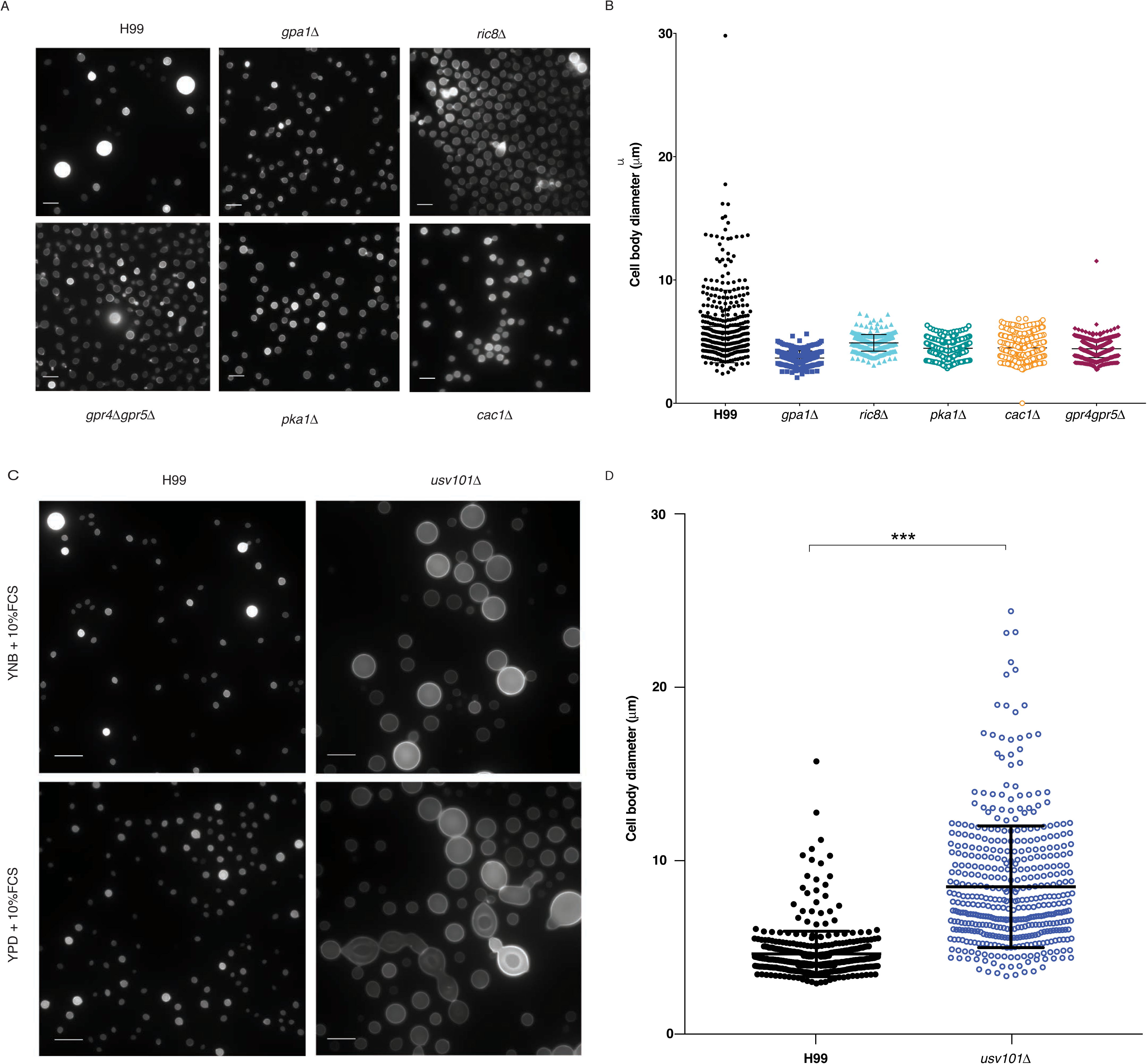
The cAMP/PKA pathway and the Usv101 transcription factor influence *in vitro* titanisation. A,B) Wild-type, *gpa1Δ, ric8Δ, gpr4Δgpr5Δ, pkalΔ*, and *cac1Δ* cells were incubated under Titan inducing conditions. A) Representative micrographs for fixed cells are shown (CFW). Scale bar = 10 μm. B) Cell size was quantified for the indicated strains. Significance was assessed using Kruskal-Wallis and Dunn's multiple comparisons (p<0.0001 for all strains relatively to H99). C) Wild-type and *usv101Δ* cells were incubated under the indicated conditions. Representative micrographs for fixed cells are shown (CFW). D) Cell size was quantified for wild type and *usv101Δ* cells induced in YNB+10% FCS. Significance was assessed using Mann Whitney U (p<0.0001).

### Usv101 negatively regulate *in vitro* Titanisation

The C_2_H_2_ transcription factor Usv101 is a master regulator of *C. neoformans* pathogenesis that negatively regulates capsule and acts downstream of Swi6, a regulator of cell cycle progression [31-33]. Usv101 is additionally predicted to regulate Gpa1 but is not itself directly influenced by cAMP. Although *USV101* has been shown to be dispensable for virulence, murine infection results in delayed dissemination to the brain, and is characterized by pneumonia rather than meningitis [31]. We therefore investigated the role of Usv101 in *in vitro* Titanisation. Compared to the H99 parent, *usv101Δ* produced significantly more titan cells *in vitro* (Fig. 4C,D; 39.25±6.45%, p<0.0001). The *usv101Δ* mutant also produced large cells when pre-incubated in YPD and then stimulated with 10% HI-FCS (Fig. 4C). Like washed wild type cells, YPD-grown *usv101Δ* cells frequently exhibited defects in cytokinesis and bud morphology. These data suggest that Usv101 may regulate nutrient-mediated priming of the yeast-to-Titan transition.

### *In vitro* Titanisation can predict *in vivo* Titanisation

We examined the capacity of non-H99 strains to produce Titan cells *in vitro*. Titanisation has not been reported for *Cryptococcus gattii*, and no increase in cell size was observed for *C. gattii* isolate R265 (Fig. S3A). Next, we screened 62 environmental and clinical *C. neoformans* isolates representing VNI, VNII, and VNB clades [34]. Strains were classified as Titanising, non-Titanising, or Indeterminate (Fig. S3B). A wide variety of cell sizes were observed in response to inducing conditions, and representative strains are shown in Figure 5A. Isolates Zc1, Zc8, and Zc12 are in the VNI clade and are capsular, thermotolerant, and melanising, comparable to H99 (Fig. 5A, S3C). Across the 62 isolates, we observed a wide range in Titanisation capacity in both clinical and environmental isolates from each clade, including clinical strains with defects in Titan cell production (Zc1, Zc12; Fig 5A), and environmental strains that produced Titan cells (S8963, Ze14 (VNB-B); Fig S3B). We also observed non-H99 clinical strains that Titanised (Zc8; Fig 5A) and environmental strains that did not (Ze18 (VNB-B), Fig S3B). Together, these data suggest that the yeast-to-Titan switch is a conserved morphogenic transition that can occur across the *C. neoformans* var. *grubii* species complex, but that individual isolates within each clade exhibit different capacities to form Titans.

**Figure 5:**
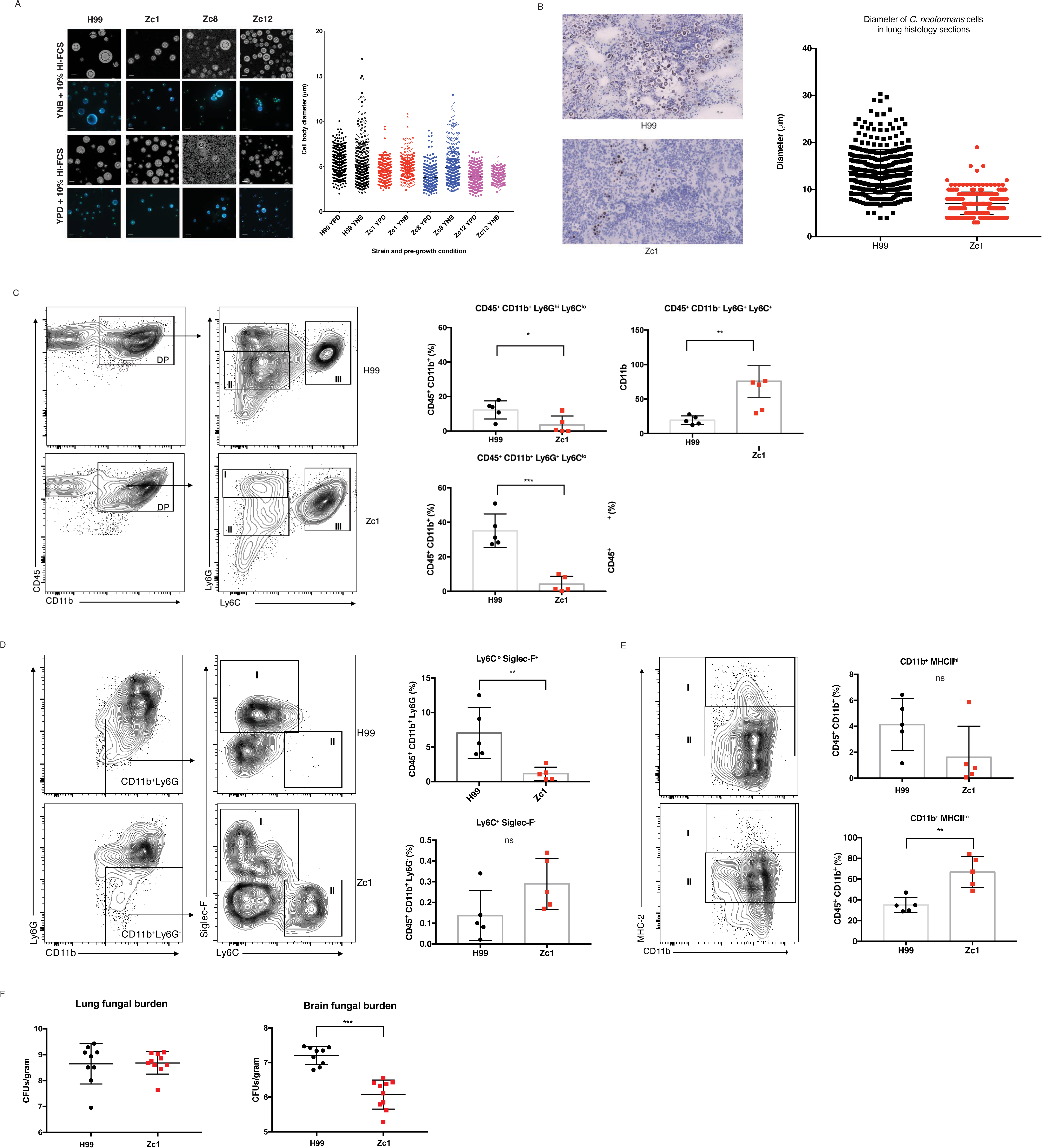
*In vitro* Titanisation predicts *in vivo* outcome. A,B) H99, Zc1, Zc8, and Zc12 clinical isolates were pre-cultured in either YPD or YNB and then inoculated OD_600_=0.001 in 10% FCS (3 days, 37°C, 5%CO_2_). A) Representative micrographs of live cells stained with CFW and SytoxGreen or counterstained with India ink and cell size quantification are shown. B) Representative lung histology from mice infected under the indicated conditions and sacrificed on day 7. Cell size was measured for each condition (n>500 cells for each condition). C,D) FACS analyses for immune cell recruitment to the lungs for mice infected with H99 or Zc1 (day 7 p.i.). C) Concatenated analysis for 5 mice under each condition were gated for CD45+ CD11b+ populations and then analysed for Ly6G and Ly6C markers. D) Indicated populations of CD45+ CD11b+ cells in individual mice were analysed for the two conditions (* p=0.0299; **p=0.0079; ***p<0.0002). F) Zc1 and H99 infected mice were followed for 21 days post-infection. Lung (p=0.907) and brain (p<0.0001) fungal burdens were measured on day 21.

Finally, we validated the capacity of our *in vitro* assay to predict *in vivo* outcome in a murine inhalation model of infection, the current gold standard for Titanisation analysis, using the type strain H99 and a clinical isolate predicted not to form Titans, Zc1 (Fig. 5A, C, p=0.0184). Mice infected with Zc1 or H99 were observed for 7 days and then sacrificed, and the lungs and brain were collected. Notably, there were clear differences in lung pathology despite comparable CFUs at this site (Fig. S3D). Lungs from H99-infected mice exhibited large lesions or granuloma in contrast to lungs from Zc1-infected mice, which exhibited fewer or no apparent lesions or granuloma (Fig. S3E). Histology also revealed foci of encapsulated fungi in the lungs of H99-infected mice, with heterogeneous cell size including both Titan (>15 μm) and yeast (<10 μm) cells (Fig. 5B). In contrast, histology of the lungs of Zc1 infected mice revealed disseminated infection, with encapsulated yeast distributed throughout the lung parenchyma. The population was more uniform in size, with the vast majority of cells less than 10 μm (Fig. 5B).

Titanisation is associated with altered immune response and increased dissemination to the brain [7, 19]. We therefore investigated the impact of the two strains on immune response in the lungs on day 7 post-infection. With the distinct nature of capsules between Titanising and non-Titanising isolates, we spectulated that these two groups might express unique PAMPs and thus differ in their interactions with immune cells. For this purpose, we focued our attention on cells of myeloid origins. The total number of CD45 cells was not significantly different (p=0.158). We observed recruitment of leukocytes into the lungs of both groups of mice, primarily comprised of CD11b+ granulocytes (Fig. 5C,D). Three distinct subsets characterized by Ly6G and Ly6C expression were observed: mature neutrophils (Ly6G^hi^, gate I) and to populations of immature Ly6G^int^ neutrophils expressing Ly6G^lo^ (gate II) and ly6C^hi^ (gate III) (Fig. 5C). While H99infected mice had more mature neutrophils (12% vs. 3.7%; p=0.0299, gate I) as well as Ly6C^lo^ immature neutrophils (37.6% vs 5.2%; p = 0.0002, gate II), Zc1-infected mice exhibited significantly higher percentage of the Ly6Chi immature neutrophil pool (Fig 5C, 76.1% vs 19.5%; p = 0.0079, gate III). The remaining CD11b+ non-neutrophil compartment contained, amongst others, eosinophils (SiglecF+) and monocytes (Ly6G^-^ Ly6C^hi^) (Fig 5D). Eosinophils were found to be higher in H99-infected mice (53.5% vs 28.8%, p=0.0085; Fig. 5D, gate I) and monocytes were elevated in Zc1-infected mice, although the difference did not reach significance (Fig. 5D, gate II; p=0.3095). We also observed differences in the frequency of cells expressing the MHC-II molecule involved in antigen presentation (Fig 5E). Although CD11b+ MHC-II^hi^ cells (gate I) were present in both groups, mice infected with H99 displayed an increasing trend of this subset, while Zc1-infected mice had a significant increase in CD11b+ MHCII^lo^ cells (p=0.0079, gate II) (Fig. 5E).

Using mouse body weight loss as a correlate of disease progression, no significant difference was observed in disease severity between Zc1 and H99 in a 21-day survival assay (p=0.0377), although Zc1-infected mice experienced a dip in weight on day 7 (Fig S3G, p=0.002). Consistent with this, lung CFUs were similar on the day of sacrifice (p=0.3411) (Fig. S3G). However, when dissemination was examined, brain CFUs on Day 21 were significantly higher in mice infected with the H99 isolate (p<0.0001; Fig. 5F). This is consistent with previous observations of differential tropism and immune response when Titan cells are present [19]. Taken together, these data suggest that our *in vitro* assay accurately predicts *in vivo* Titanisation and that isolates that form Titan cells drive quantitatively distinct immune responses from those that do not.

## Discussion

The yeast-to-Titan transition is a host-specific morphogenic switch that can influence disease outcome. Titan daughters have altered stress resistance compared to their mother cells, and the presence of Titans is associated with altered immune status [6-8]. Despite their importance, mechanisms underlying Titanisation have been challenging to dissect due to the lack of a reproducible *in vitro* induction protocol. Here we present a rapid, robust *in vitro* induction protocol that generates cells with all the properties of *in vivo* Titan cells, recapitulates previously identified regulators (Gpr4/Gpr5), directly confirms the role of cAMP pathway elements (Gpa1, Cac1, Pka1) and identifies new regulators (Ric8, Usv101). The assay further accurately predicted an *in vivo* defect in Titanisation by a clinical isolate with otherwise intact virulence.

We define *in vitro* Titans based on cell body size, and show that in our assay approximately 15% of H99 cells form Titans within three days. Other reports have included capsule size when measuring Titan frequency, with a cut-off of 15 μm, and our data agree with the observation that capsule is influenced by cell cycle [35]. However, our data also support the observation that Titan capsule is more compact than yeast phase capsule [18, 24], and the relationship between capsule size and cell body size changes as cells cross the 10 μm threshold. For example, for the cells shown in Figure 1A, the yeast cell is 4.98 μm with an 18.49 μm capsule (ratio of 3.70), while the Titan cell is 21.9 μm with a 37.17 μm capsule (ratio of 1.69). We therefore suggest that measurements that include capsule may inaccurately quantify Titan frequency by including yeast cells with large capsules and excluding smaller Titans with dense capsule. For this reason, we have limited our analysis to cell body size, defining Titans as those cells greater than 10 μm.

There are some differences between our findings and the published literature. First, we identify a subpopulation of cells with very thin cell walls and altered cell shape, which we term Titanides. These cells appear to be distinct from previously described microcells (<1μm) and yeast daughter cells. Based on their altered cell wall, Titanides are likely to differ in the exposure of host relevant ligands relative to yeast cells. Future work will further characterize these cells and investigate the role of this sub-population in pathogenesis.

Second, single daughter cells isolated from Titan mothers were diploid or, in some cases tetraploid, whereas colonies arising from Titan mothers tended to be comprised of primarily haploid or aneuploid cells (Fig 2B). This is in contrast to daughter cells dissected from *in vivo* Titan mothers, which tended to be aneuploid or haploid[6]. This finding highlights an important aspect of Titan cell biology that has proved challenging to study. Titan cells are thought to allow phenotypic diversity through the generation of aneuploid daughters with increased access to the fitness landscape as a result of changing gene dosage[6]. Current models suggest that uninucleate, highly polyploid mothers bud off haploid or aneuploid daughters, requiring asymmetric DNA division via an unknown mechanism. Efforts to understand the molecular mechanisms underlying this process will benefit from an *in vitro* model, and our data already suggest that the diversity of these daughter cells is greater than previously described.

Finally, the clinical isolate Zc1 and the clinical type strain H99 elicited distinct immune responses. While both H99 and Zc1 strains induced leukocyte recruitment into the lungs, granulocytes predominated the response and these could be clustered into three unique subsets (a mature neutrophil subset (Ly6G^hi^) and two distinct immature neutrophil subsets (Ly6C^lo^ and Ly6C^hi^)). H99 infection was associated with increased frequency of mature neutrophils and the LyC6^lo^ immature neutrophil subset whereas the Ly6C^hi^ immature neutrophils were dominant during infection with the Zc1 isolate. It is not known whether these cells express different effector functions and what their polarization state is and thus more work is required to better understand if they mediate protection or susceptibility to infection with *C. neoformans*. Other noble differences in immune responses involved disparate frequencies of eosinophils and monocytes recruited during infection. Increased frequency of eosinophils and CD11b^+^MHC-II^hi^ was observed in mice infected with H99 relative to the Zc1 isolate while the frequency of monocytes and CD11b+MHC-II^l^° cells was higher in Zc1-infected mice. Overall, it is currently unclear where these differences in immune status drive different disease outcomes. Regardless, Zc1-infected mice exhibited moderate lung pathology and reduced dissemination to the brain suggesting that there might be qualitative differences immune responses driven by Zc1 vs H99. While the overall survival of mice infected with H99 or Zc1 was comparable, our data suggests that morbidity might be an important factor to consider during infection with different isolates. Indeed, it is possible that other differences between the two strains are driving these differences in pathogenesis. Zc1, like H99, is a VN1 clade patient isolate with no defects in capsule, thermotolerance, or melaninsation. Titanisation capacity appears to be the single largest difference between Zc1 and H99, and we identify a wide range in Titanisation phenotypes for environmental and clinical isolates. The relative impact of Titanisation on pathogenesis and clinical outcomes is a pressing question and future work will investigate this further, particularly in the context of immune-altered states such as neutropenia, and T and B lymphocytopenia.

Our *in vitro* model offers some initial insights into the underlying molecular mechanisms regulating Titanisation *in vivo*. First, *in vitro* Titans form following a Prime-Induce signal. *C. neoformans* growth in minimal media is known to alter the expression of secreted proteins relative to YPD and influences stress resistance through transcriptional and post-translational changes [36, 37]. Exposure to serum is a known signal for capsule induction [38]. However, the combinatorial impact of these conditions on *C. neoformans* morphogenesis had not yet been tested. Importantly, unlike capsule induction, which is nearly 100% penetrant across the population, *in vitro* Titans make up a minority of the total population, suggesting that transcriptional or post-translational priming interacts with the micro-environment and/or specific state of individual cells.

Second, previous work has strongly suggested a role for the cAMP signal transduction cascade in Titanisation *in vivo* [20, 27]. Because *gpa1Δ, pka1Δ*, and *cac1Δ* strains are rapidly cleared from the host lung, direct testing of their role in Titanisation was not possible *in vivo* [20, 28, 29].

Here, we confirm this model through direct demonstration that cells deficient in adenylyl cyclase activity fail to form Titans *in vitro*. However, despite the requirement for adenylyl cyclase activity, constitutive activation is not sufficient to induce Titans. The addition of exogenous dcAMP does not induce Titanisation nor does it restore Titanisation to cAMP/Pka pathway mutants in our *in vitro* assay. Consistent with this, hyperactivation of the pathway using GAL-inducible *PKA1* and *PKR1* constructs produces a heterogeneous population of both large, polyploid cells and yeast phase cells after 48hr [22]. Similarly, expression of a constitutively active version of the Gα protein Gpa1 doubles the percent Titan cells *in vivo* [20]. In each of these cases, Titan cells make up a fraction of the total cell population. These data suggest that Titan induction via cAMP/Pka1 interacts with individual cell state to determine cell fate.

Third, we identify the transcription factor Usv101 as a negative regulator of Titanisation. Cells lacking *USV101* form Titan cells at a high frequency compared to the parental strain and also form Titan-like cells when induced from YPD pre-cultures. When induced from YPD, both wild-type and *usv101Δ* cells tended to accumulate defects in cytokinesis, which was not observed in YNB-pre-cultured Titans, suggesting that nutrient-pre-exposure primes cells for Titan-phase budding. Nutrient limitation alters the expression of density-dependent factors [36], and our data suggest that nutrient pre-conditions may also influence subsequent morphological transitions.

The lack of an *in vitro* protocol has hampered efforts to study the specific morphogenic processes distinguishing Titan and yeast-phase growth. However, differences in nuclear segregation and bud development programs are consistent with previous observations of highly polyploid uninucleate mother cells generating aneuploid daughter cells via asymmetric nuclear division. Usv101 is predicted to act in parallel to the cAMP pathway and downstream of the cell cycle regulator Swi6. The interaction between cell cycle and pathogenicity factor expression is an emerging theme in *C. neoformans* biology: Recent work has also highlighted cell cycle regulation of pathogenicity factors [33] and the cyclin Cln1 has been shown to regulate capsule and melanin, both of which are regulated by cAMP and negatively regulated by Usv101 [31, 32, 35, 39].

Finally, we relate the *in vitro* induction protocol to physiologically relevant *in vivo* conditions. *C. neoformans* is a globally distributed environmental fungus. The mammalian lung is not thought to be a reservoir for *C. neoformans*. Rather, our data suggest that the host lung may serve as a niche for bacterial-fungal interactions that mediate pathogenesis. Titan cells are observed in the host lung and BAL can replace FCS as the inducing compound for YNB-primed cells. Antibiotic treatment to reduce bacterial lung burdens reduced Titan induction in a murine inhalation model, and co-culture of live or heat-killed *E. coli* with *C. neoformans* led to Titanisation. Exposure of primed cells to NMAiGn, a synthetic version of the bacterial cell wall component MTP, was sufficient to induce Titan cells. MTP and its derivatives are known to activate cAMP-mediated morphological transitions in *Candida albicans* and other ascomycetes[27]. Together, these data point to a conserved mechanism for bacterial-fungal interactions underlying pathogenic morphological transitions.

## Methods

### Media and strains

Strains used in this study are summarized in Table 1. *C. neoformans* H99 [40], *gpa1Δ, cac1Δ[28]*, and *pka1Δ*[41] were gifts from Andrew Alspaugh, Duke University, NC, USA. The gpr4Δ gpr5Δ [42] was kindly provided by Joseph Heitman, Duke University, NC, USA. Strains *ric8Δ* and *usv101Δ* were obtained from the Madhani 2015 collection (NIH funding, R01AI100272) from the Fungal Genetics Stock Centre and were validated by PCR and shown to phenotypically match published strains [30, 31]. *C. gattii* R265 [43] was provided by Neil Gow, University of Aberdeen, UK. Isolates are summarized in Table S1. Cells were routinely cultured on YPD (1% yeast extract, 2% bacto-peptone, 2% glucose, 2% bacto-agar) plates stored at 4°C. For routine culture, cells were incubated overnight in 5 mL YPD at 30°C, 150 rpm. For Titan induction, cells were incubated overnight at 30°C, 150 rpm in 5 mL YNB without amino acids (Sigma Y1250) prepared according to the manufacturer’s instructions plus 2% glucose. Fetal Calf Serum (FCS) was obtained from either BioSera (Ringmer, UK) or Sigma, which both induced Titan cells to a similar degree. FCS was routinely stored in 5 ml aliquots at ‐20 to prevent repeated freeze-thaw cycles. FCS was heat-inactivated by incubation at 56°C for 30 min.

### Titan cell measurement

Cells were induced and either fixed with 4% methanol free paraformaldehyde and permeabilised using 0.05% PBS Triton-X and stained for total chitin with calcofluor white (CFW, 10 μg/ml) and DNA with SytoxGreen (Molecular Probes, 5 μg/ml) or stained live using calcofluor white (CFW, 5 μg/ml) and the cell permeable nucleic acid stain SybrGreen (Invitrogen, 0.5X). SybrGreen preferentially stains dsDNA, but has low affinity for ssDNA and ssRNA, so is not appropriate for quantitative DNA analysis. However, it does allow visualization of nucleic acid dynamics within live cells. Cells were imaged using a Zeiss M1 imager for fixed cells and either a Zeiss Axio Imager or a Nikon Eclipse TI live imager, both equipped with temperature and CO_2_ control chambers for live imaging. To visualize capsule, live cells were counterstained using India Ink (Remel; RMLR21518). Representative images are shown. Cell diameter was measured using FIJI, with frames randomly selected, all cells in a given frame analysed, and at least three images acquired per sample for each of two independent runs, representing experimental duplicates. In all instances, n>200 cells. Statistical analyses were performed using Graphpad Prism v7. For pairwise comparisons, the Mann-Whitney test was applied. For multiple comparisons, ANOVA and Kruskal-Wallis were applied.

### Flow Cytometry for cell ploidy

Cells were fixed and stained according to the protocol of Okagaki et al 2010[3]. Briefly, cells were fixed with 4% methanol free paraformaldehyde and permeabilised using 0.05% PBS Triton-X. Cells were washed 3 times with 1x PBS and stained with 300 ng/ml DAPI. Where indicated, samples were enriched for large cells by passing through an 11 μm filter prior to staining. Cells were analysed for DNA content using an LSRII flow cytometer on the Indo-1

Violet channel and 10,000 cells were acquired for each sample. Data were analysed using FlowJo v. 10.1r7. Doublets and clumps were excluded using the recommended gating strategy of SSC-H vs SSC-W followed by FSC-H vs. FSC-w, and cells were then gated to exclude autofluorescence using unstained pooled haploid and diploid control cells. The gating strategy is provided in Fig S1B. Gates for 1C, 2C, 4C, and 8C were established using H99 (haploid) and KN994B7#16 (diploid) controls incubated in rich media conditions as described above.

### Co-culture with *E. coli*

H99 cells were incubated overnight at 30°C, 150 rpm in 5 ml YNB without amino acids + 2% glucose. Cells were inoculated into 1xPBS+ 0.04% glucose (the concentration of glucose present in serum) in the presence or absence of 10^7^ live or heat-killed *E. coli*. Co-cultures were incubated for 48 hr at 37°C 5%CO_2_ and assessed for Titan formation by microscopy as described above.

### Compound identification

To identify compounds of interest, total HI-FCS (Biosera) was loaded onto an AKTA purifier system from GE Healthcare with an Agilent column: Bio SEC-3, 100A, 4.6x300mm at a flow rate of 0.3 ml/min for size exclusion chromatography. For the initial run, 400 μl serum was run in phosphate buffer (25 mM NaH2PO4, 150 mM NaCl, 0.01% NaN3, 2 mM EDTA, pH 7.2), collecting 100 μl fractions in a 96 well plate. The entire plate was screened for capacity to induce Titans by incubated H99 cells pre-grown in YNB+Glucose at OD_600_=0.01 in 10% fraction+1xPBS in a 96 well plate format. Plates were examined for Titan cells after 48 and 96 hr. The entire assay was run in triplicate and twice independently using distinct lots of FCS. Inducing fractions were pooled and lyophilized. The residue was resuspended in MeOH and desalted. Then, the solution was submitted to HPLC separations, which were carried out using a Phenomenex reversed-phase (C_18_, 250 × 10 mm, L × i.d.) column connected to an Agilent 1200 series binary pump and monitored using an Agilent photodiode array detector. Detection was carried out at 220, 254, 280 and 350 nm. The entire volume was purified by RP-HPLC using a gradient of MeOH in H_2_O as eluent (50–100% over 70 min, 100% for 20 min) at a flow rate of 1 ml/min. The main fraction was dried, suspended in a minimal volume of DMSO-d6 and submitted to ^1^H-NMR, ^1^H-^13^C HSQC and ^1^H-DOSY analyses. NMR data were acquired on a Bruker 500 MHz spectrometer.

### Mice and infection models

All animal experiments were performed under UK Home Office project license PPL 70/9027 granted to DMM in accordance with Home Office ethical guidelines. C57BL/6J mice were bred and maintained in individually ventilated cages (IVCs) at the Medical Research Facility at the University of Aberdeen. For each experiment, group size was determined based on previous experiments as the minimum number of mice needed to detect statistical significance (p<0.05) with 90% power (α=0.05, two-sided). Mice were randomly assigned to groups by an investigator not involved in the analysis and the fungal inocula were randomly allocated to groups. Inocula were delivered in a blinded fashion. Mice were provided with food and water *ad libitum*. Mice were monitored for signs and symptoms of disease. Weight was recorded daily. Mice showing weight loss of greater than 30% and signs of disease progression were immediately culled by a schedule one method (cervical dislocation).

C57BL/6J male mice (n=5/group, 8-12 weeks old) were anesthetized using injectable anesthesia and infected intranasally with 20 μl PBS suspension containing 10^5^ *C. neoformans* (H99 or Zc1) pre-grown in Sabouraud Dextrose medium [8]. For 7 day studies, mice were culled by euthatal injection. Lungs and brains were collected under sterile conditions. Whole brains and one lung lobe were weighed and homogenized for CFU counts. For long term infection study, mice were observed for a least 21 days with daily records of mice body weights as one of the correlates of disease progression. Mice were culled at the end of the experiment by cervical dislocation. Survival data were assessed by Kaplan Myer and Gehan-Breslow-Wilcoxon test.

### Lung Flow Cytometry

Lungs from infected mice were used to generate single-cell suspension using mouse lung dissociation kit and the gentleMACS as per manufactures’ instructions (Miltenyi Biotec). Fungal cells were separated from mammalian cells via a 70%/30& discontinuous Percoll gradient centrifugation. Immune cells were stained with the fixable viability dye eFluor 455UV (eBiosecience) for 30 min at 4 °C, washed with 1x PBS then fixed with a 2 % paraformaldehyde (PFA) solution for 10 min at room temperature. Cell surface staining with antibody cocktail of mAbs specific to CD45-BV650, MHC-II-PECSF594, CD11c-APC, CD11b-BUV395, Ly6C-PE, Ly6G-FITC (all from BD Biosciences) was performed in FACS buffer containing 2 % fetal calf serum, 2 mM sodium azide and anti-CD16/32 for 30 min at 4 °C, washed then acquired on the BD Fortessa cell analyser (BD Biosciences). FlowJo software v10 (Tree Star) was used for data analysis.

### Statistical Analysis of FACS data

Data represent percent live CD45+ cells. Statistical analyses were performed using Graph Pad Prism (v 7), and significance was determined using Mann-Whitney U test. Bars represent 95% CI. Variance within the groups was not statistically different (F test to compare variances).

### Histology and *In vivo* Titanisation

For lung histology, C57BL/6J male mice (n=5/group, 8-12 weeks old) were anesthetized and infected intranasally with 20 μl PBS suspension containing 10^5^ *C. neoformans* (H99 or Zc1) pregrown in Sabouraud Dextrose as above. After 7 days, mice were culled by euthatal injection. Lung sections were preserved in OTC medium and sectioned (2-4 μm) for histology. Fungi were visualized by silver staining and hematoxylin counterstain (Sigma HHS32) using the Sigma-Aldrich Silver Stain modified GMS kit according to the manufacture’s instructions).

### Bronchoalveolar lavage (BAL)

BAL was performed on male C57BL/6 mice (8-12 weeks) culled by CO_2_ exposure. Lungs were perfused with 1 ml 1xPBS and the collected fluid concentrated by overnight drying on a speedvac. The resulting pellet was weighed, resuspended in sterile PBS, and used at a concentration of 10% w/v in place of FCS in the induction protocol. Animal experiments were performed by ID, ERB, AC, and DMM.

## Acknowledgements

We thank Prof Joseph Heitman and Prof Kirsten Nielsen for helpful discussions. We are grateful to Attila Bebes and Linda Duncan in the Iain Fraser Cytometry Centre (Aberdeen University), and Debbie Wilkinson, Lucinda Wight and Kevin MacKenzie in the Microscopy and Histology

Core Facility (Aberdeen University) for their expert help with the cytometry and microscopy experiments. We are also grateful for the assistance of staff at the University of Aberdeen Medical Research Facility.

## Author contributions

ID, RY, GDB, DMM, and ERB planned the experiments. ID, TD, AC, LTS, NL, RY, DMM, and ERB performed the experiments. ID, RY, and ERB performed the flow cytometry analyses. TR, MF, TB, TH, MJ, RM, and GDB provided materials and valuable insight during this study. ID and ERB drafted the manuscript with contributions from TD, AC, LTS, NL, TR, MF, TB, TH, MJ, RM, GDB, RY, and DMM.

---

**Figure S1:**
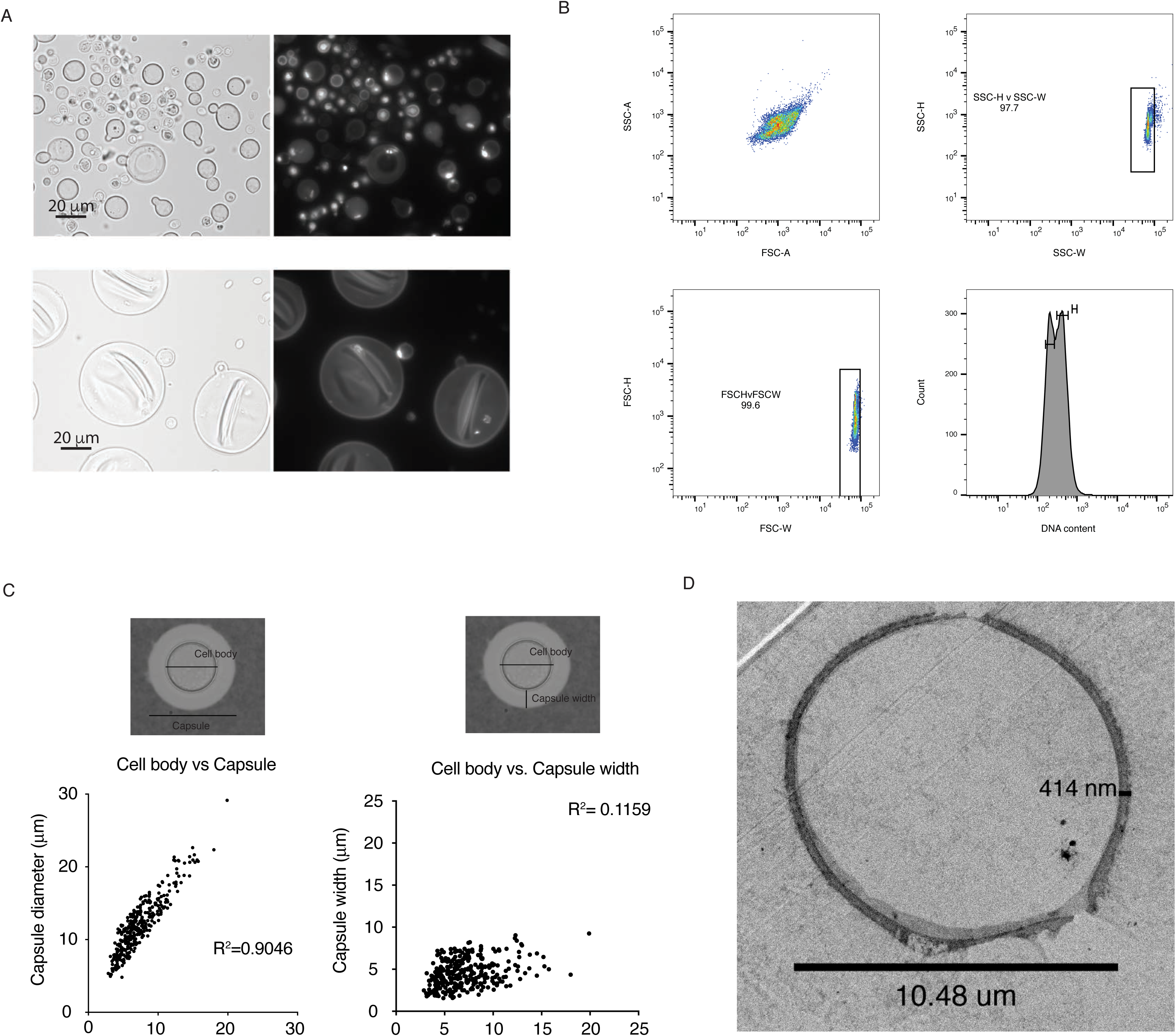
Characterisation of Titan cells. A) H99 was pre-grown in YNB + Glucose overnight at 150 RPM, 30°C and inoculated at OD_600_=0.5 and 0.01 into 10% FCS. Representative cells are shown after 7 days. Cells were fixed, permeabilised, and stained (DAPI, 300 ng/ml) to visualize nuclei. Scale bar = 20 μm. Imaged on a Zeiss Axio Observer Z1 at 40x. B) Cell body and capsule diameter were measured as indicated for H99 cells induced using the established protocol (n=299). Capsule width was defined as capsule – cell body diameter. R^2^ values are shown. C) TEM of a Titan cell showing cytoplasm excluded by a large vacuole.

**Figure S2:**
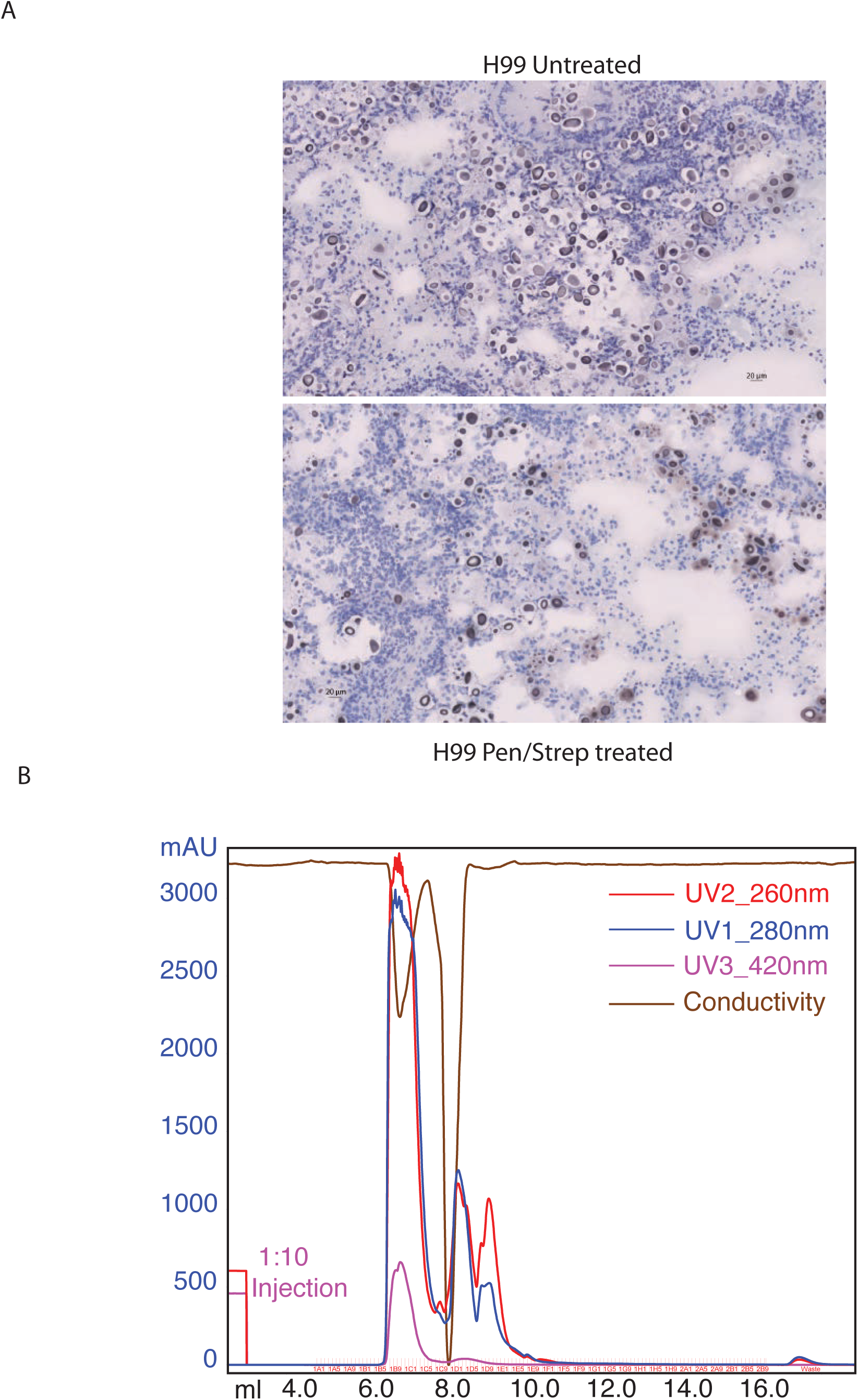
Serum fractionation by SEC. Total HI-FCS was fractionated by size exclusion chromatography. The chromatogram of the total volume is shown.

**Figure S3:**
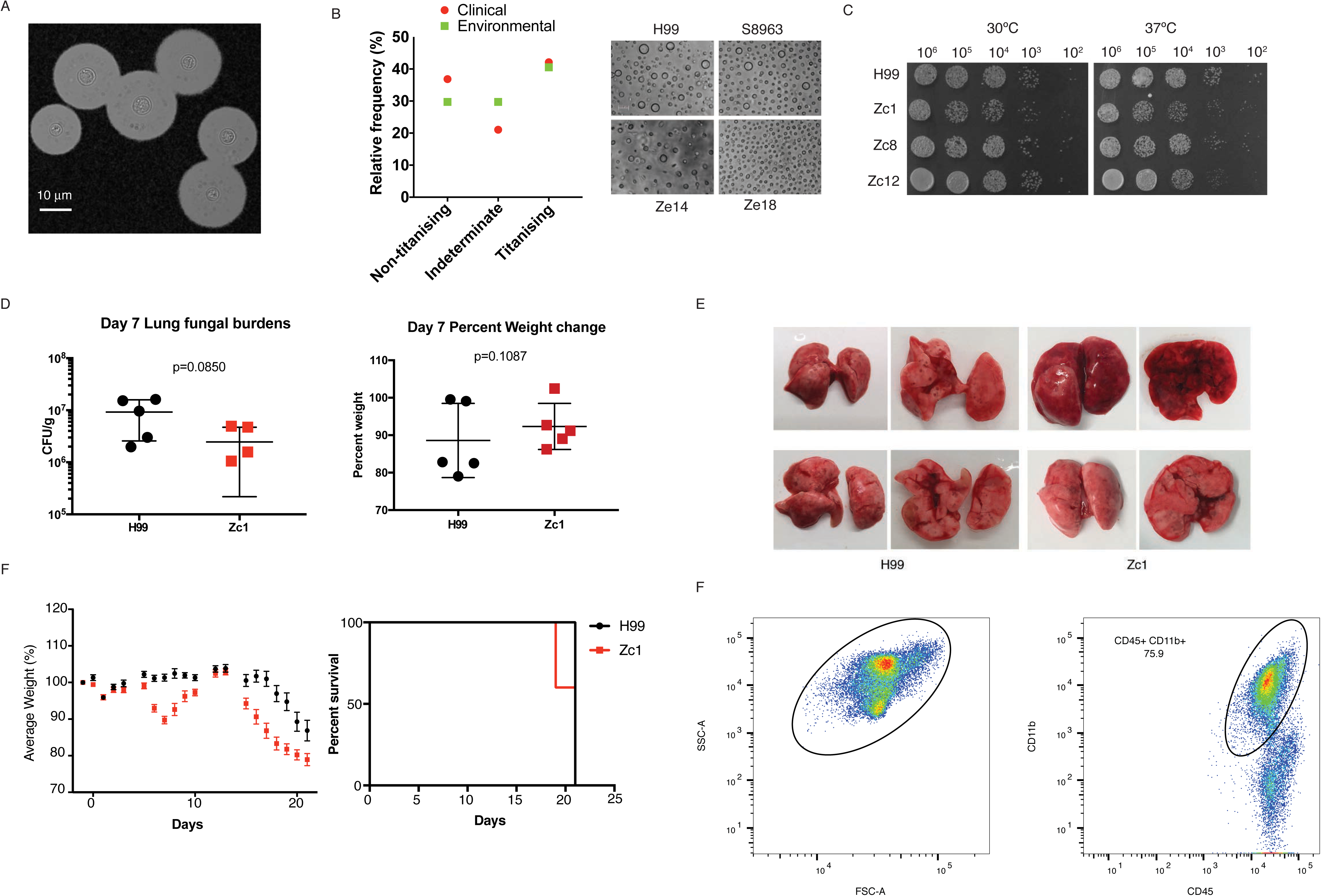
Titanisation and *in vivo* clinical and environmental isolates. A) Cryptococcus gattii strain R265 was pre-grown in YNB, inoculated into 10%FCS at OD=0.01, and incubated at 37°C, 5%CO2 for 5 days. Cells were counterstained with India ink to reveal capsule. Scale bar = 10 μm. B) 63 Clinical and environmental isolates were induced for Titan cells (YNB, 10%FCS, OD600=0.001) and analysed for increased cell size and cell ploidy (DAPI, flow cytometry). Strains were categorized as Titanising if cells >10 μm were observed, indeterminate if cells >7μm but < 10μm were observed, and non-Titanising if only cells < 7um were observed. The percent of strains identified for each category is shown. Representative environmental isolates S8963, Ze14, and Ze18 are shown compared to H99. C) Clinical isolates Zc1, Zc8, and Zc12 were grown in YPD and then spotted on to YPD agar and incubated at 30 or 37°C as indicated. D, E) Mice infected with H99 or Zc1 for 7 days were sacrificed and D) the lung fungal burden and percent weight loss recorded. E) Images of representative lungs from infected mice. G) Disease severity was monitored for 21 days by weight loss (Mann-Whitney U, p=0.002) and mice were sacrificed at humane end-point (p=0.0377). F) Gating strategy for immune cell recruitment in the lungs of infected mice.

**Table S1:**
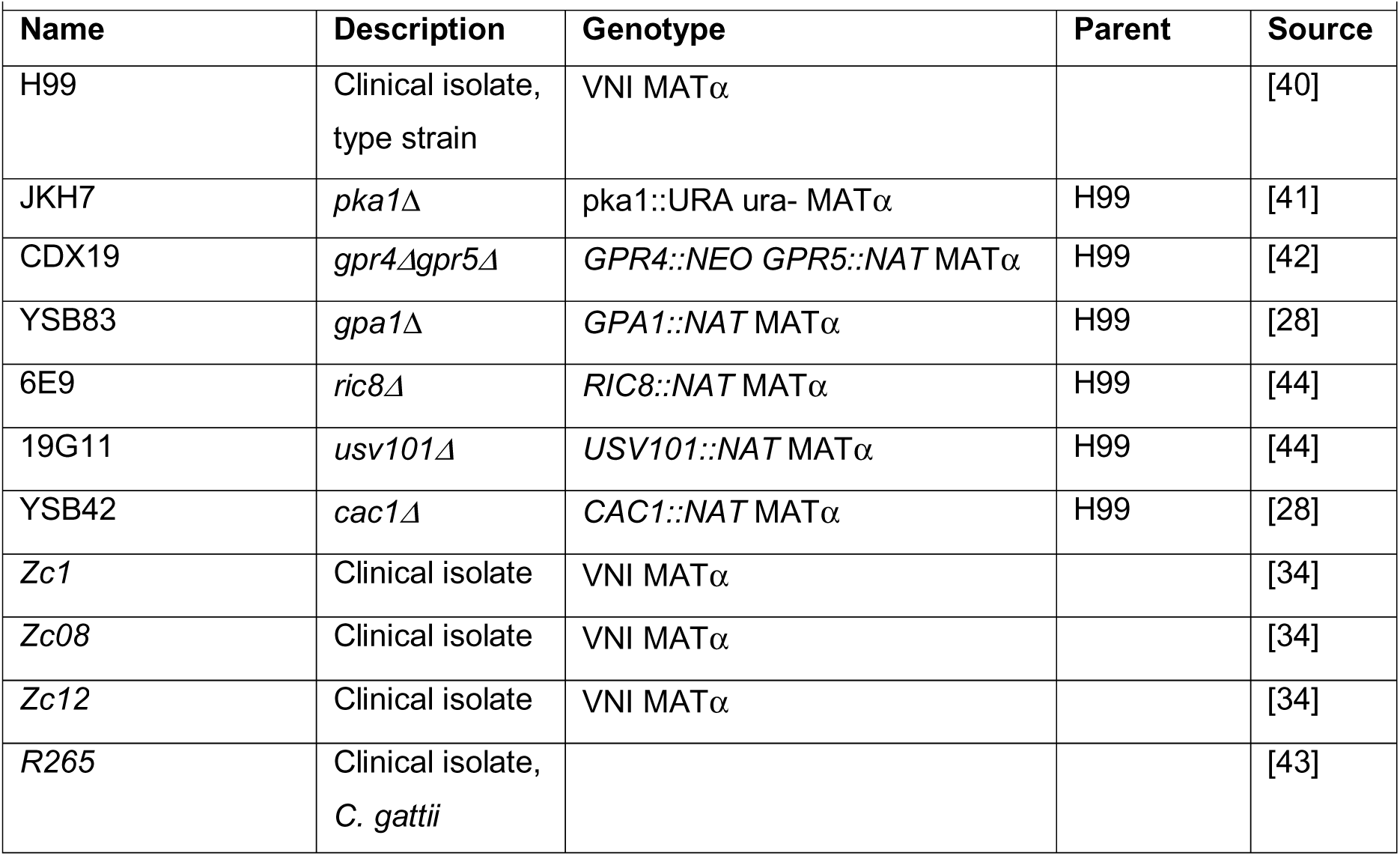
Strains used in this study?

